# Urban wastewater contains a functional human antibody repertoire of mucosal origin

**DOI:** 10.1101/2024.06.02.597066

**Authors:** Sean Stephenson, Walaa Eid, Chandler Hayyin Wong, Elisabeth Mercier, Patrick M. D’Aoust, Md Pervez Kabir, Stefan Baral, Kimberly A. Gilbride, Claire Oswald, Sharon E. Straus, Alex Mackenzie, Robert Delatolla, Tyson E. Graber

## Abstract

Wastewater-based surveillance of human disease offers timely insights to public health, helping to mitigate infectious disease outbreaks and decrease downstream morbidity and mortality. These systems rely on nucleic acid amplification tests for monitoring disease trends, while antibody-based seroprevalence surveys gauge community immunity. However, serological surveys are resource-intensive and subject to potentially long lead times and sampling bias. We identified and characterized a human antibody repertoire, predominantly secretory IgA, isolated from a central wastewater treatment plant and building-scale wastewater collection points. These antibodies partition to the solids fraction and retain immunoaffinity for SARS-CoV-2 and Influenza A virus antigens. This stable pool could enable real-time tracking of correlates of vaccination, infection, and immunity, aiding in establishing population-level thresholds for immune protection and assessing the efficacy of future vaccine campaigns, particularly those that are designed to induce humoral mucosal immunity.

## INTRODUCTION

The COVID-19 pandemic has driven a worldwide interest in wastewater-based surveillance (WBS). The development of assays to quantify viral nucleic acid has been rapidly adopted by public health organizations around the world. Applied in the context of WBS, the sensitive, and target-specific quantitation of viral nucleic acid originating from infected host excreta enables the detection of a range of pathogens, and this measured signal serves as a proxy for disease incidence, followed in near-real time as it waxes and wanes in the targeted community (1–7).

The surveillance initiatives with the greatest impact to-date have focused on the detection of SARS-CoV-2 RNA extracted from sewage samples collected at local water resource recovery facilities (WRRFs; representing sewered households) or other strategic up-sewer facilities such as airports, long-term care facilities, and hospitals (8). Analogous methods have been used to successfully detect and quantify other viral RNAs such as Respiratory Syncytial Virus (RSV), and Influenza A/B (IAV/IBV) (9–13). Recently, these methods have also been rapidly adapted and applied to monitor Mpox virus DNA (14–16). These proxy measures of disease provide actionable intelligence for public health stakeholders, and when interpreted through publicly accessible web portals, empower individuals and communities alike to take effective action to reduce the burden of disease.

While current WBS platforms can accurately track trends in disease incidence, they cannot directly quantify the degree of immune protection of a monitored community facing an actual or potential emerging pathogen. Traditional methods to do so include serological measurement of pathogen-specific antibodies, epidemiological data analysis, vaccination coverage assessment, as well as longitudinal cohort studies. Combining these approaches helps public health officials gauge immunity and guide control and prevention strategies effectively.

Antibodies or immunoglobulins (Ig) are large glycoproteins of the vertebrate adaptive immune system that help identify and classify foreign biomolecules (i.e., antigens) in the body through direct binding. Serum concentrations of specific subsets of antibodies correlate with onset of infection and/or a vaccination event and have the potential to protect from subsequent infection and/or symptomatic disease (i.e., correlate of protection; COP). Seroprevalence surveys provide valuable information about the degree of immune protection in populations, and in the case of SARS-CoV-2 can distinguish past natural- vs. vaccine-mediated exposure (17). However, their requirement for significant testing infrastructure and often long turnaround time (weeks-to-months) can limit their usefulness to public health teams (e.g., informing decisions about the timing of a vaccination campaign) especially in under-resourced contexts. In view of these impediments, we explore here if measurement of wastewater antibody levels represents a potentially valuable addition to the immune surveillance armamentarium.

The demonstration of antibodies in wastewater is a potential first step towards tracking community immunity levels, opening the possibility of defining immune protection thresholds for a given pathogen and monitoring the efficacy of a mucosal vaccine campaign. Protein mass spectrometry has identified significant amounts of human Ig chain peptides in wastewater (18) and a preprint by Agan et al (19) found that wastewater collected at both building- and WRRF-scales contained SARS-CoV-2 specific IgG and IgA antibodies by enzyme-linked immunosorbent assay (ELISA). However, the extent of intact and functional human antibody pools in urban wastewaters remains unclear.

There are five Ig classes present in human serum: IgA, IgG, IgM, IgD, and IgE, that can be distinguished by their heavy polypeptide chains. The typical antibody titre response triggered by respiratory infections such as SARS-CoV-2, RSV, and IAV has been thoroughly characterized (20–22). In general, IgM is the first antibody to respond, detectable within a week of infection, peaking at 2-3 weeks, while IgA, a key component of mucosal immunity, appears in the respiratory tract and serum within the first 1-2 weeks. IgG is detectable between 2-3 weeks and remains detectable months after infection or vaccination (23,24). These observations are typically monitored in serum, and less commonly in saliva/sputum (25,26). The levels of these Ig classes generally correlate with clinical severity in the case of COVID-19 (27,28). Mechanistically, immune protection of SARS-CoV-2 infection and symptomatic COVID-19 is thought to be primarily through neutralizing serum IgG and IgA antibodies (29).

In considering the potential use of municipal wastewater to monitor human antibodies, it’s important to recognize that relative and absolute isotype abundances in sputum, saliva and feces which contain antibodies secreted from mucosal surfaces, are radically different from that in serum. In serum of a healthy individual, IgG levels are approximately ten times greater than IgA and IgM (30). Although high titres in serum offer the best sensitivity, other sample types such as sputum, urine, or stool serve as less invasive, but ostensibly less sensitive, ways of monitoring the immune response of an individual to pathogens like SARS-CoV-2 (31–33). In one study, feces and serum from healthy controls contained comparable amounts of IgA (per unit of input), while there were two orders of magnitude less IgG in feces vs. serum (34). Another study reported approximately 10-fold more IgA vs. IgG titres in sputum from healthy volunteers (35). Comparatively, urinary immunoglobulin concentrations in healthy individuals are vanishingly small -- several orders of magnitude lower than feces per unit of input for both IgG and IgA (36). IgM (30), IgE, and IgD (37) are comparatively minor isotypes in all these contexts but should not be discounted *a priori* as a detectable COP.

Mucosal surfaces, as the main interface with the environment, serve as potential entry points for pathogens. In addition to cell-mediated protection, the adaptive immune system relies predominantly on secreted IgA to protect these areas (38). Given that the aggregate gastrointestinal, respiratory and genitourinary tract mucosal surface area in a healthy adult is 30-40 square meters, perhaps unsurprisingly, over 80% of Ig-producing cells in the body are found in the mucosal immune system (39). The major secreted Ig, dimeric IgA (or secretory IgA; SIgA), attaches to pathogens blocking their interaction with host cells preventing tissue penetration and propagation (40). Similarly, but with notable differences, IgM can also be secreted and has a natural affinity for viruses, bacteria, and fungi (41). IgM monomers can form pentamer and hexamer structures that aid with activation of the complement-dependent cytotoxic pathways and clearing of pathogens at sites of infection (42). Polymerization of both IgA or IgM monomers and secretion of the polymeric antibodies from B-cells is facilitated by Joining chain (JC), a small scaffold protein, highly expressed in exocrine tissue, that bridges neighbouring heavy chains through disulphide bonds. Once secreted from B-cells, the JC of polymeric antibodies interacts with the polymeric immunoglobulin receptor (pIgR) on the basolateral surface of, for example intestinal epithelial cells, triggering transcytosis into the intestinal lumen, where a fragment of pIgR, secretory component (SC), remains associated with the secreted antibody complex, serving to enhance both stability and function (43).

We demonstrate here that bulk protein content partitions to wastewater solids. Using a combination of western blotting, ELISA, and mass spectrometry, we identify a robust repertoire of intact human antibodies, predominantly secreted IgA (SIgA). Further, we identify wastewater-derived IgA and IgG populations with affinity/avidity for SARS-CoV-2 S protein as well as Influenza A virus antigens. Detectable, albeit weak signals were present in building-scale samples where people infected with SARS-CoV-2 resided, while WRRF-scale samples showed consistent detection of both SARS-CoV-2 S IgA and IgG antibodies during a major vaccination campaign that coincided with a large epidemic wave in Fall 2023.

## RESULTS

### Bulk protein content partitions to wastewater solids

Compared to cellular monocultures or tissue lysates, wastewater samples are highly heterogenous. We and others noted early in the COVID-19 pandemic that SARS-CoV-2 RNA-associated particles partition to the solids fraction of wastewaters (2,44,45). The partitioning phenotype of a nucleic acid target is influenced by the nature of the material (e.g., viral or another encapsulated particle or condensate) from which it is extracted. Proteins would be expected to behave similarly, with partitioning dictated by the structures they are extracted from, or in the case of isolated proteins such as antibodies, their innate biophysical properties. However, protein partitioning in different wastewater types (WRRF vs. building-level) and fractions (solids vs. liquid) has not been extensively characterized.

We first investigated how protein partitions into solids and liquid fractions in primary sludge and post-grit influent samples obtained from the Ottawa WRRF (Post-grit influent refers to the wastewater stream after removal of grit, dense particles like sand and gravel, which is followed by the primary treatment, where primary sludge is formed from the settling of solids in clarifiers). The solids fraction is mainly insoluble material pelleted at high-speed centrifugation, while the resulting supernatant/liquid contains more soluble components. Processing the solids fraction requires homogenization to solubilize proteins, while the liquid fraction requires concentration of the proportionally larger volume to yield significant amounts of protein (**Fig 1A**). We chose a homogenization buffer containing PBS and 0.5% Triton X-100 to maintain near-physiological pH and to retain the native state of proteins while helping solubilization (46). For concentration of liquid fractions, ultrafiltration with a molecular weight cut-off of 10 kDa was chosen to retain as many soluble proteins as possible.

**Figure 1.**
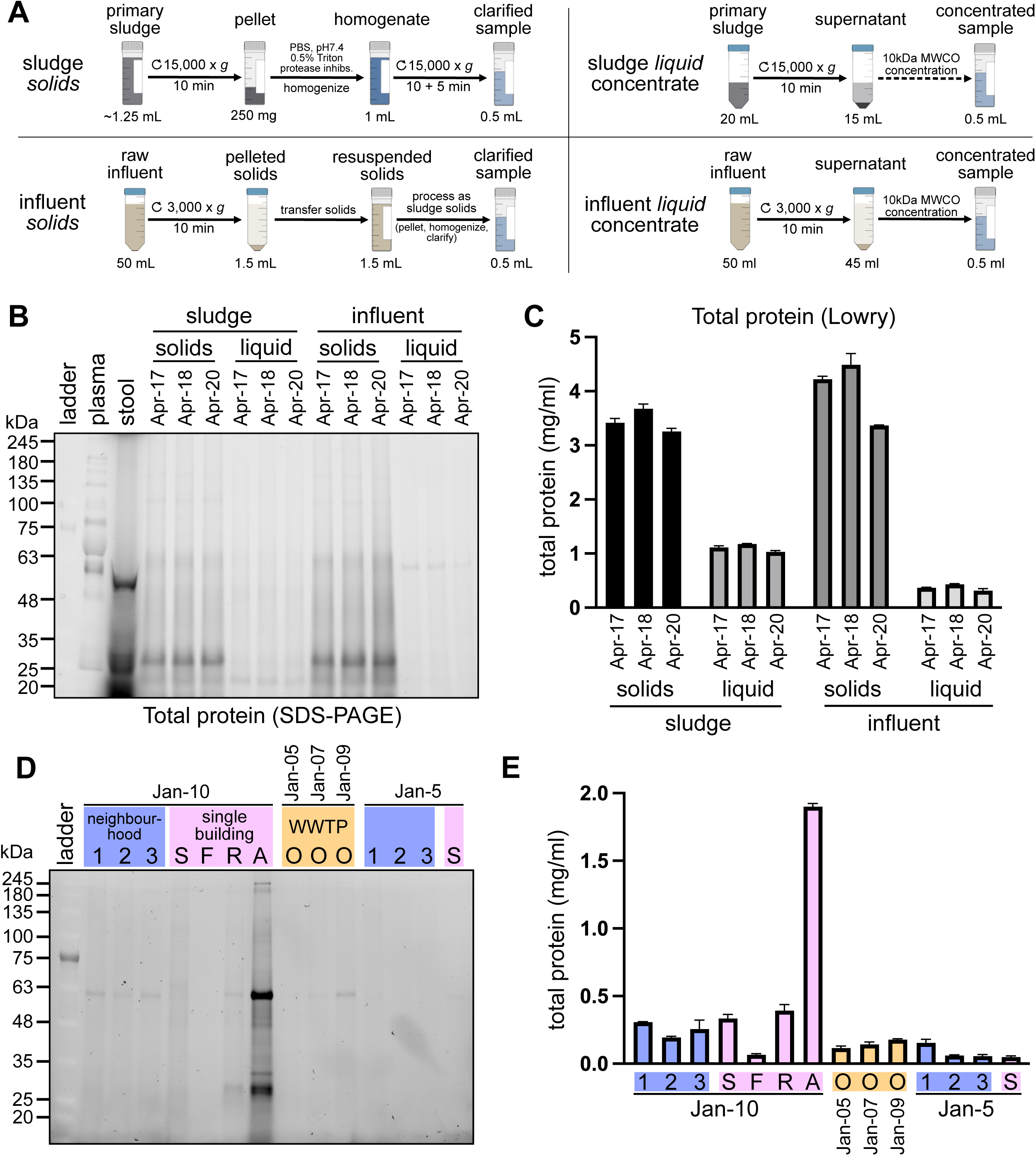
Total protein profiles assessed in wastewaters by Stain-Free (tryptophan) labelling. (**A**) Schematic of clarification and concentration methods used to process WRRF influent, primary sludge, or in-sewer wastewater (processed in the same manner as “raw influent”) for protein analyses. (**B**) Stain-Free SDS-PAGE profiles of protein extracts (equal volumes) from 24h-composited primary sludge and influent solids, as well as the liquid concentrate fractions collected over 3 different days in April 2023 at the Ottawa WRRF. Plasma and stool obtained from healthy volunteers were used as comparators to assess integrity and concentration of wastewater proteins. (**C**) Total protein extracts from B were quantified using a colorimetric Lowry-based assay with BSA as a reference standard. Protein concentration is expressed as mg of total protein per ml of assay input. Error bars represent standard deviation of technical triplicates. (**D**) As in B, but with equal volumes of total influent obtained from building-, neighbourhood- and city-scale sampling sites in January 2023. Three different neighbourhoods (#1-3), 4 buildings (identified by single-letter codes), and 1 central treatment plant (Ottawa; O) were profiled. (**E**) As in C, but with samples from D.

We assessed total protein content of extracts using stain-free PAGE and measured yields using a BCA assay with BSA as a reference standard. In paired primary sludge and influent samples collected from the WRRF on three separate days, a similar profile was observed in the extracts from the solids and concentrated liquid fractions of each sample (**Fig 1B**). Quantitation of total protein from the same samples replicated the qualitative findings (**Fig 1C**), showing that roughly two-thirds of the bulk protein content partitions to the solids fraction in primary sludge, while for influent solids, this proportion is approximately 90%. Consistent with the finding that the bulk of protein is associated with suspended solids, screening of equal volumes of influent obtained from building-, neighbourhood-, and city/WRRF-sampling sites show very low protein content albeit with a wide range of levels observed between samples, demonstrating the need to normalize any measured target to allow comparisons between sites and across time (**Fig 1D, E**).

### Secretory IgA makes up the majority of an antibody repertoire detected in wastewater solids

Having shown the bulk of protein partitions to primary sludge and influent solid fractions, we next screened these for human IgG and IgA by western blot. A primary ∼55kDa band, the expected size of their heavy chains, was consistently observed in both anti-human IgG and IgA blots in multiple samples from different dates (**Fig 2A**). Consistent with our bulk protein content findings, little if any difference was observed in the electrophoretic mobility or amount of the two Ig heavy chain classes between equal amounts of primary sludge or influent solids loaded. Neither IgA nor IgG heavy chains were observed in the primary sludge and influent liquid concentrates by western (**Fig S1**). We also observed heavy bands consistent with IgM chains by western (**Fig S2**).

**Figure 2.**
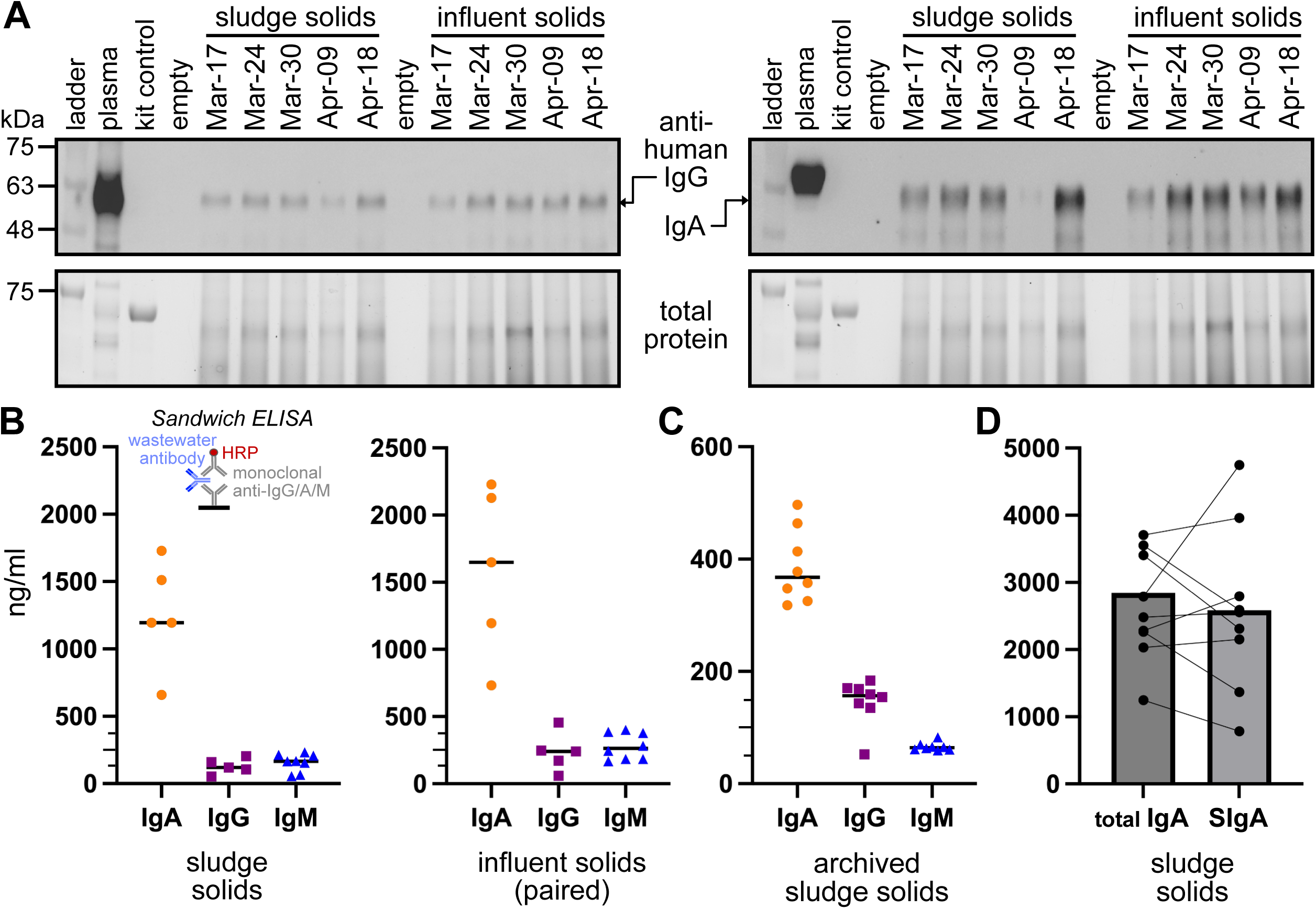
Measurable quantities of intact total human Ig isotype pools in wastewater fractions. (**A**) Western blots (reducing conditions) of primary sludge or influent solids probed for human IgG (left) or IgA (right). Human plasma was included as a positive control. Stain-free images of the gel are shown prior to blotting and indicate total protein loaded (bottom). (**B**) Sandwich ELISAs quantifying Ig classes in different sample types and time periods; equal volumes of processed, freshly obtained, paired primary sludge (left; n=5 different days collected spring 2023), and influent solids (right). (**C**) As in B but with archived/frozen primary sludge solids (right, n=8) from a prior time period (late 2022) were assayed. (**D**) Same sample comparison of total IgA and SIgA sandwich ELISAs of primary sludge solids (n=8) from a 3^rd^ time period. Quantities are based on a standard curve and are expressed as ng of Ig per ml of assay input.

To confirm intact Ig classes in wastewater, we next measured human IgA, IgG, and IgM by quantitative ELISA. Freshly processed solids (Spring 2023) from influent and date-paired primary sludge samples (n=5-8 days) were measured. Total Ig was readily detected within the dynamic range of the assay, with significantly higher human IgA levels compared with IgG or IgM observed. Similar relative amounts between primary sludge and influent solids from paired (collected the same day) samples were recorded (**Fig 2B**). Ig classes in frozen/archived primary sludge samples from a different collection period, in late 2022 (n=8 days) were also measured. As observed in the fresh solids, the 3 Ig classes were above the assays’ limits of quantification in the 8 frozen samples tested (**Fig 2C**). We noted that although IgA still predominated, both it, and IgM made up a smaller fraction of the Ig classes than in the fresh samples collected in Spring 2023.

The high concentration of IgA in wastewater relative to IgG is not unexpected since the bulk of the human excreta contributing to the wastewater pool would be expected to be non-plasma, instead originating from mucosal surfaces (e.g. feces, saliva, nasal mucus). With this in mind, we measured the proportion of SIgA from the total IgA antibody pool from primary sludge solids sampled in Winter 2024 by ELISA. Importantly, SIgA accounted for most of the total IgA signal (**Fig 2D**).

### Mass spectrometry confirms solids partitioning, isotypes and secretory antibody complex proteins

As a complementary approach to ELISA, mass spectrometry (MS) was used to profile peptides from wastewater fractions. WRRF samples were again resolved by reducing SDS-PAGE, then gel regions close to expected antibody light (L) and heavy (H) chain sizes were excised and subjected to tandem LC-MS (**Fig 3A**). Consistent with prior experiments, peptides that mapped to antibody chains were found to predominate in primary sludge and influent solids, rather than the liquid fractions (note that no peptides could be mapped for the influent liquid fraction). Moreover, α heavy chains (IGHA1) were mapped at the highest intensity, followed by consistent but weak detection of μ chains (IGHM) and only a single peptide mapping to γ chain (IGHG) (**Fig 3B, and summarized in Fig 3C**). Neither δ nor ε chains were detected, suggesting that IgD and IgE did not make up a significant fraction of the wastewater antibody repertoire at the time of collection.

**Figure 3.**
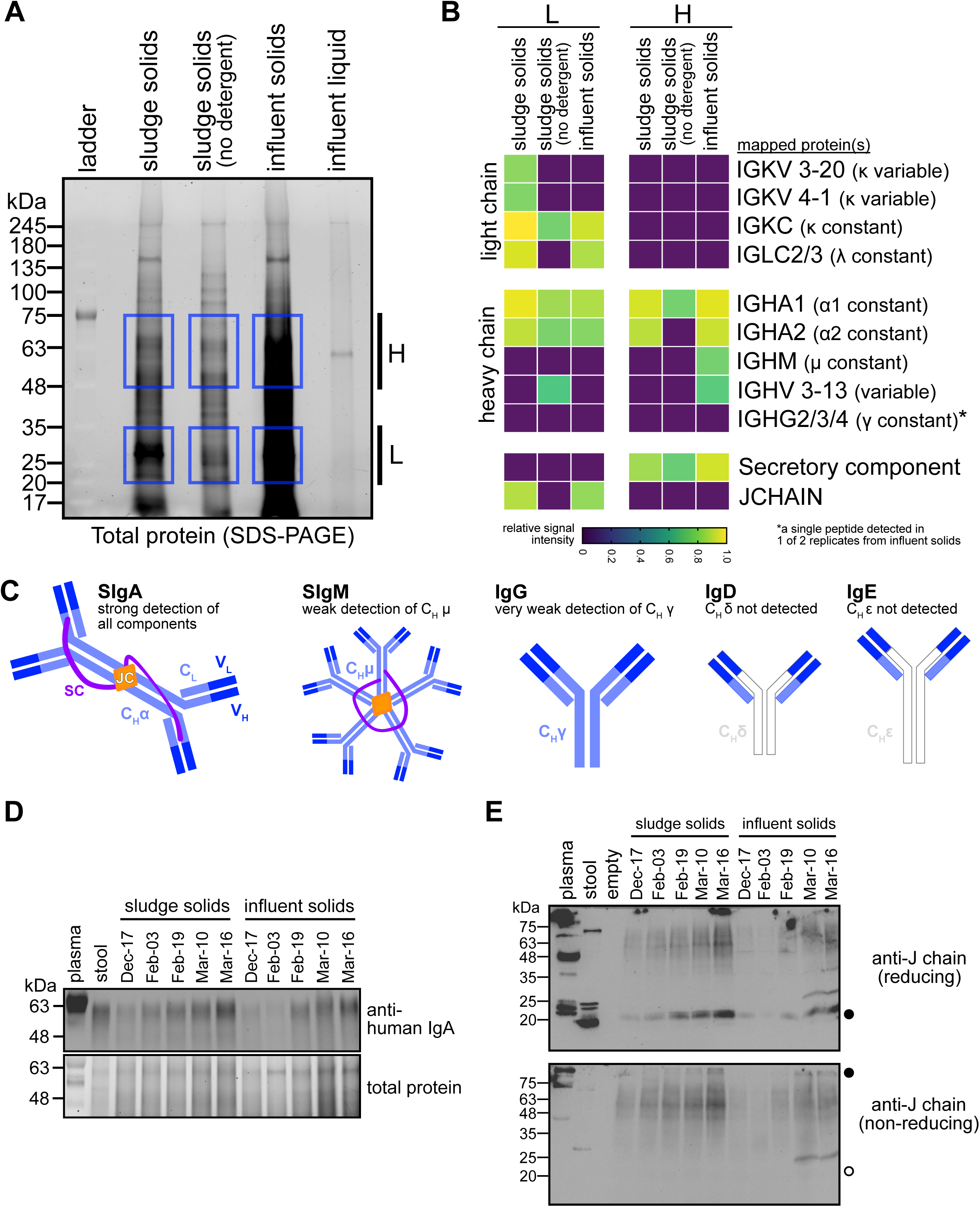
Ig, JC, and SC polypeptides are present in wastewater with a significant fraction of JC associated with high molecular weight complexes. (**A**) Total protein gel (reducing) of wastewater samples showing regions of interest (H; high molecular weight, and L; low molecular weight) submitted for analysis by MS. Note the relatively weak protein profile of influent liquid concentrate which was not analyzed further. Sludge solids that were homogenized by dilution in the absence of detergent and protease inhibitors were also submitted for MS analysis (sludge solids - no detergent). (**B**) Heatmap of peptide intensities across regions of interest and sample type. Rows are grouped by light, heavy, and mucosal Ig chains and accessory proteins, the latter of which uniquely associate with secreted antibody sub-classes. (**C**) Pictogram summary of the MS findings. (**D**) Additional wastewater samples collected winter 2022/2023, together with human plasma and stool samples were probed to confirm the presence of IgA. (**E**) These same samples were analyzed for the presence of the ∼18kDa JC under reducing conditions (top panel, closed circle) and JC-associated high-molecular weight complexes (closed circle) under non-reducing conditions (bottom panel). Open circle marker indicates the location of JC signal observed in wastewater samples under reducing conditions, that shifts to putative Ig complexes in non-reducing conditions.

Of note, we observed significant peptide intensities mapping to *heavy* chains that migrated at an apparent molecular weight of ∼20-35 kDa (gel region “L”). This suggests significant partial degradation and/or proteolysis of heavy chains. On the other hand, the solids fractions revealed high intensity detection of κ and λ light chain peptides and variable regions were also mapped suggesting the possibility that epitope binding could be preserved.

We also detected strong intensities of peptides mapping to SC and JC. Both defining components of SIgA, they are secreted and amongst the most highly expressed proteins in the gut and respiratory system ((47); proteinatlas.org). The SIgA ELISA data suggested that wastewater-derived SC is at least partially associated with intact dimeric IgA. To determine if JC is similarly associated with Ig in wastewater, we turned to non-reducing western blots, which preserves disulphide bridges between light and heavy chains as well as in polymeric IgA and IgM. Human IgA was robustly detected in wastewater samples collected in Winter 2023/2024 and we noted that the mobility of IgA in wastewater was like that in human stool, suggesting comparable glycosylation profiles, and thus body compartment(s) of origin (**Fig 3D**). High amounts of JC were readily detected in reducing conditions by western blot, including in plasma and stool (where slower-moving, putatively glycosylated species, could also be observed; (48)), at the expected size (**Fig 3E**, top, marked with closed circle). Importantly, in non-reducing conditions, we observed high molecular weight bands consistent with JC associated with Ig complexes of mucosal origin (**Fig 3E**, bottom, marked with closed circle). Free JC was absent in these conditions (open circle marker in **Fig 3E**, bottom) suggesting that the wastewater-derived JC fraction is entirely complexed. Together, these data indicate that a SIgA pool is present in wastewater solids and that it is likely dimeric, associated with SC and JC.

### Antigen-binding activity is preserved in the wastewater antibody repertoire

We next asked whether the sampled antibody repertoire contained intact *and* functional Igs with preserved antigen-binding activity -- a prerequisite for monitoring specific correlates of disease and immune protection therefrom. Respiratory pathogen targets associated with existing wastewater, clinical surveillance and public health response plans were first considered, namely, SARS-CoV-2, Respiratory Syncytial virus, and Influenza A/B virus. In the Fall of 2023, a population seroprevalence survey in the province of Ontario showed that >85% of blood donors were positive for SARS-CoV-2 anti-N IgG, while 100% were positive for anti-S IgG (https://www.covid19immunitytaskforce.ca/seroprevalence-in-canada/), distinguishing infection-acquired and vaccine-induced immunity, respectively. Concomitantly, a vaccination campaign with formulations targeting the XBB.1.5 omicron variant had begun, and one of the highest incidence periods of SARS-CoV-2 infection in Ottawa had not yet peaked. Therefore, anticipating high levels of SARS-CoV-2 IgG and IgA, we chose samples collected and archived from the Ottawa WRRF from this period and tested for the presence of SARS-CoV-2 S-specific antibodies by semi-quantitative IgA or IgG ELISAs that have specificity for the trimeric ectodomain of ancestral/Wuhan S protein (**Fig 4A**). A stool sample donated from a COVID-19 hospital in-patient (collected ca. 2020) and diluted 1:10 with buffer served as a specificity control. This indicated titratable IgA, but no IgG signal, consistent with a strong mucosal response to a natural infection (**Fig 4B**).

**Figure 4.**
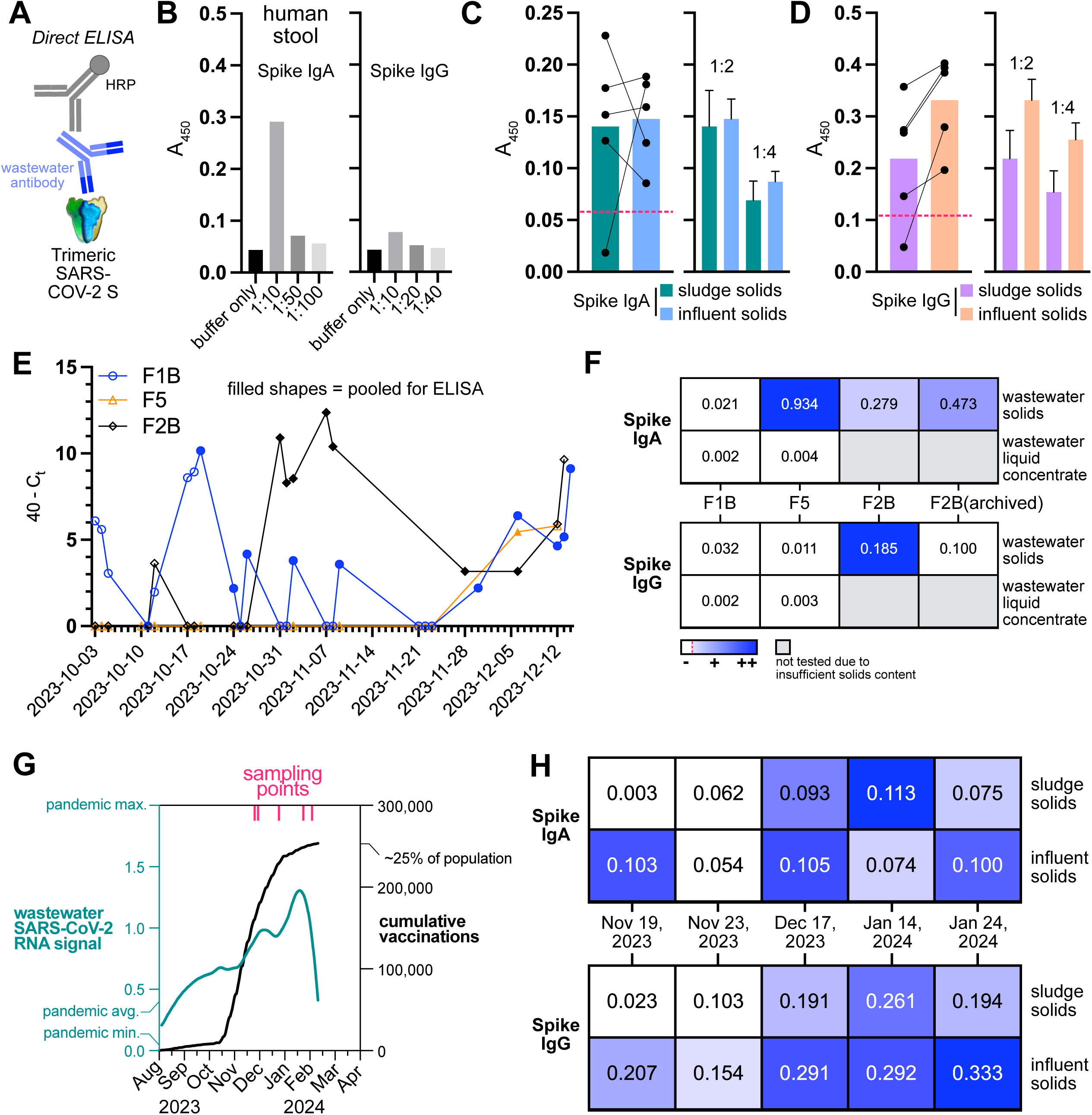
Wastewater contains IgA and IgG pools with preserved antigen-binding function that correlate with disease incidence and vaccine uptake in the sampled communities. (**A**) Schematic of the anti-SARS-CoV-2 IgA/G direct ELISAs used. The assays specifically detect antibodies to trimeric SARS-CoV-2 Spike protein (Wuhan). (**B**) Stool isolated from a COVID-19 patient shows titratable Spike IgA, but very low IgG quantities demonstrating expected mucosal isotype expression. (**C)** Spike IgA expressed as blank-subtracted absorbance values (450 nm) in both primary sludge and influent solids sampled from different days (paired samples, n=5; left) exhibited linearity-of-dilution (right). Dotted line indicates the baseline reading of kit negative control. (**D**) As in C, but specific for Spike IgG (n=5). (**E**) SARS-CoV-2 N1 RT-qPCR positivity across a sampling period at an emergency housing facility. (**F**) SARS-CoV-2 anti-IgA and IgG detection in building-scale wastewater pooled from E. (**G**) Wastewater samples presented in C, D were collected during a period of high SARS-CoV-2 incidence and vaccination as indicated by high wastewater RNA signal (LOESS model of daily measurements) and high community uptake of bivalent vaccines. (**H**) Temporal trends in anti-Spike protein IgA (top) and IgG (bottom) during this period in samples collected at Ottawa WRRF.

Next, we tested date-paired influent and primary sludge solids (n=5) from the WRRF collected over a 10-week period. 4 of 5 samples collected from the WRRF and diluted 1:2 in buffer to minimize matrix effects were positive for both anti-trimeric Spike IgA (**Fig 4C**) and IgG (**Fig 4D**), based on signal above the kit negative control. Consistent with total protein and Ig pools, there was little if any difference noted between primary sludge and influent solids for IgA, while there was a trend of more Spike-specific IgG in influent solids.

While WRRF-scale detection is important for monitoring overall trends in antibody levels, assessment of building-scale trends is an attractive use-case scenario wherein the proximity in time and space to excretion into wastewater might favour more intact and therefore measurable antibody pools. We chose to assess wastewater from 3 emergency housing facilities (providing accommodation to people experiencing homelessness) located in the City of Toronto during Fall 2023, a time when, similarly to Ottawa, the population was experiencing the 2^nd^ largest pandemic wave and a vaccination campaign. We monitored wastewater-based SARS-CoV-2 RNA activity in the buildings using RT-qPCR, detecting the N1 locus (49). We then pooled samples that were N1-positive and contained enough (>50 mg) of solids to ensure the highest possible Ig content. Samples were concentrated, and equal volumes were assessed with the same S-specific IgA and IgG ELISAs as above. For the solids fractions, high-positive Spike IgA was found in 2 of 3 facilities (F1B, F5) while Spike IgG positivity (low) was found in only 1 of the 3 facilities. F2B wastewater, that had been frozen and thawed was re-assayed to determine stability of the signal in archived samples. These data show that building-scale detection of disease-specific antibodies is possible from processed wastewater solids.

We attempted to relate WRRF/city-level trends in SARS-CoV-2 IgA/G with time-resolved RNA-based measurements and vaccine uptake using the samples measured in Figure 4C, D. These samples were collected during the waxing and peak times of a very high pandemic wave (∼70% of the weekly pandemic peak wastewater SARS-CoV-2 RNA signal; **Fig 4G**) that coincided with a strong bivalent vaccine uptake in the community (cumulative doses representing 25% of the population). Indeed, although the signal magnitudes were low, IgA (which does not see a strong increase upon vaccination, but may do so with infection, or re-infection) signal appeared to remain stable during the sampling period (**Fig 4H**, top heat map). In contrast, there was an increasing trend observed for IgG (whose expression increases with vaccination) during the same sampling period.

To extend the application to other antigens, we measured anti-IAV and -RSV IgA in Ottawa WRRF and Toronto shelters during the winter of 2022 when both viruses were causing an epidemic and in the Spring of 2023 approximately one month following the IAV and RSV waves. There was very low, but detectable anti-IAV and anti-RSV IgA at Ottawa WRRF throughout the two collection periods. Intriguingly, several of the Toronto shelter samples showed clear detects for anti-IAV IgA (**Fig S3**). Together, these data indicate the pool of antibodies in wastewater retains immunoaffinity for human antigens.

## DISCUSSION

### Partitioning of bulk protein in wastewater

Both primary sludge and influent have different protein profiles within their respective solid and liquid fractions, which can be observed by the presence or absence of distinct bands at a given molecular weight after separation by SDS-PAGE. However, primary sludge and influent solids fractions are noticeably similar compared to the liquid fractions. Some species are seen in the influent liquid fraction that are not present in the primary sludge liquid fraction; indicating that degradation, differential partitioning based on biophysical properties, and/or de novo bacterial protein synthesis or other newly introduced proteins occurs as wastewater travels through the treatment plant. This observation is intuitive, as the integrity of proteins in wastewater will deteriorate with more time spent going through the process. Physical, chemical, and microorganism-driven protease activities will affect the degree to which a protein will stay in its native state. Analogous to what has been demonstrated with DNA and RNA, proteins are likely to be partially protected by being encapsulated in complex dissolved organic matter and extracellular polymeric substances (50–52). Post-translational modifications including glycosylation could also contribute to protection from (photo)chemical and enzymatic degradation. Such particulate matter aggregates over time and is easily collected by sedimentation. Thus, it is included in all our solid fractions, which coincidently gave higher total protein yields after processing and became the focus of our downstream characterization of the constituent antibody repertoire.

Wastewater collected within the sewershed (i.e., at building- and neighbourhood-scale sewer collection points) was like influent collected at the WRRF; these samples, compared with primary sludge, are very dilute with respect to protein concentration, and the insoluble solids content is comparatively highly variable. Due to this we were only able to assess the liquid fraction in samples that had significant solids content. Total protein was highly variable between liquid fractions from different sites and likely depends on factors such as daily occupancy, the variable use of water, sampling methods and timing specific to each collection point. Samples taken further upstream from the WRRF provide a unique epidemiological benefit potentially capturing a smaller population of interest such as a community with known vulnerabilities, campus, or hospital. However, in such cases large volumes of wastewater are required to reliably detect protein targets such as antibodies in the liquid fractions. Further innovation in process engineering is needed; possibly borrowing from the knowledge gained with concentrating low-copy SARS-CoV-2 RNA from large volumes (53).

### Proportion of Ig classes in wastewater

The IgA isotype was detected at the highest proportion across all sample types and fractions relative to IgG and IgM. This trend held true for previously frozen/archived vs. freshly processed samples from two different time periods, although the overall signal for both total IgA and IgM, but not IgG, was disproportionately lower suggesting that each has its own unique stability dynamics. Two, non-mutually exclusive factors could account for the difference observed between the two time periods; the freezing/archiving may have reduced antibody affinity and avidity, or there was truly less IgA in late 2022 vs. spring 2023. We did not formally test the effect of freezing and/or long-term storage on antibody detection. However, it is interesting to note that IgG concentrations were similar between fresh and archived samples, perhaps suggesting a real reduction in IgA levels in late 2022 vs. spring 2023. Because the archived samples were collected in an earlier period than the fresh samples, it’s possible that daily/weekly/monthly distributions of total immunoglobulins were qualitatively different. Longitudinal assessment of freezing/archiving is needed to directly assess the relative stability of antibody isotypes in wastewater solids.

Using ELISA, we determined that SIgA is responsible for most of the total IgA signal observed in wastewater solids as no significant difference were observed between sIgA and total IgA levels in paired primary sludge samples. These observations correlate with what is expected in urban wastewater as major expected contributions of human secretions such as intestinal fluid, feces, and saliva are high in SIgA. IgM can also be secretory, but its seroprevalence is much less abundant and shorter-lived than IgA *in vivo* and likely explains why its signal was proportionally lower (27). Alternatively, IgG is not highly abundant in secretions, but minor amounts may still accumulate in wastewater due to contributions from non-secreted bodily fluids and with its long-lasting seroprevalence.

### Species specificity and tissue of origin of immunoglobulins in wastewater

It cannot be ignored that wastewater is a highly complex and dynamic sample type. It contains proteinaceous material from a diverse complement of prokaryotes and eukaryotes. This is especially a concern when probing for protein targets based on immunoaffinity, a notoriously promiscuous biophysical property. The presence of chemicals like detergents originating from domestic or industrial uses can bind to proteins and affect ELISA sensitivity and specificity (54). Additionally, some protein epitopes from non-human species, including those on Igs can cross-react in immunoaffinity-based experiments like ELISA or Western blot (55). Furthermore, complex samples such as human plasma suffer from matrix effects in immunoassays like ELISA (56). Such effects produce inaccurate readings due to interference with non-specific sample components or high concentrations of targeted analyte (i.e, Hook effect) and express as a non-linear dilution series. An example of this is with the quantification of cytokines, which are in relatively low concentration compared to other compounds in plasma and can be masked by matrix effects but can be more reliably analyzed with proper buffer selection and careful dilution (57,58). The dilution series that were run with our wastewater samples were not free from potential matrix effects, and therefore may affect the absolute quantification of Igs. However, samples were routinely diluted over 100-fold in commercial assay buffer when quantifying total IgA, IgG, and IgM to minimize matrix effects.

To address potential issues of specificity, we performed complementary western blot and MS experiments to support the presence of human Igs and the relative amounts of isotypes. We observed expected sizes of heavy chains for both IgA and IgG, and additionally the JC by western blot. All components of SIgA were also detected by MS: the α heavy chain, isotype-agnostic light chains, JC, and SC. These findings agree with MS-based peptide analyses on total urban wastewater fractions from Spanish municipalities, where κ light chains and the constant heavy chain of IgA were among the most abundant human proteins identified (18,59). It was unexpected that no heavy chains corresponding to IgG were detectable in our MS dataset, in contrast to the presence of both IgA and IgM heavy chains. IgG and IgM were proportionately lower than IgA when analyzed by ELISA and may not have been in sufficient quantity for reliable detection by MS. Two IgA subclasses, namely IgA1 and IgA2 were found to be equally represented in wastewater solids. This observation alone speaks to both the species and tissues of origin, as humans (along with non-human primates) are the only species known to express both. Further, the ∼1:1 proportion would be expected from mucosal tissues, in contrast to the ∼9:1 ratio of IgA1:IgA2 observed in serum (60).

Another interesting observation in the MS dataset was the apparent degradation of both the IgA1 and IgA2 heavy chains, that appeared somewhat in equal proportions in both the low and high molecular weight gel regions. This speaks to the complexity of the sample due to its pooled nature (over one million individuals and multiple tissues of origin represented), as different protein isotypes and allotypes are likely present, with a variety of post-translational modifications (e.g., glycosylation), proteolytic and/or oxidative degradation states; all of which can affect quantification. Taken together, we conclude that the values provided for total Igs are estimates of class/isotype proportion.

### On the need for normalization of inputs

An accurate temporal Ig trend requires careful normalization of measurements to correct for inter-sample variability. Here we presented Ig concentrations normalized to sample input by mass for solid fractions, and volume for liquid fractions. Critically, we noticed that samples with the same volume or mass input did not always yield similar amounts of total protein. Moving forward, we suggest using a total protein metric (e.g., derived using a BCA assay) as an easily accessible normalizer for total as well as disease-specific Igs. This could be further improved by normalizing to multiple human gene products that are both consistently observed in wastewater and correlate with human excreta contribution to wastewaters (e.g., as described in (61)). This would require a multiplexed immunoaffinity platform and adds to the specificity issues discussed above. Currently, there is a need for more research to identify good protein targets and combinations thereof that are markers of human contribution to wastewater and best capture the protein variation across different catchment sizes and seasonal changes.

### Improving sensitivity of disease-specific antibody assays for the wastewater context

Igs, particularly SIgA, appear to be in sufficient quantities to detect consistently regardless of sample date, in both primary sludge and influent solids. However, the nature of an antibody is to have a very specific target, and although we can detect total Igs in wastewater, being able to detect disease-specific antibodies such as those against SARS-CoV-2 are of true epidemiological interest. We qualitatively detected IgA and IgG antibodies that recognize the trimeric spike protein of SARS-CoV-2 in both primary sludge and influent solids. Due to the detectable, but low signal, lack of reference curve for these assays, and above-mentioned concerns about normalization, we were conservative in inferring any temporal changes/trends in signal. Disease-specific antibodies will only be a subset of the total Igs shed by an individual and will require enrichment to reliably quantify. We chose an antibody detection system with specificity for the ectodomain of ancestral trimeric S protein to ensure the highest specificity for SARS-CoV-2 at the potential risk of reduced analytical sensitivity. Indeed, the latter quality may have been further eroded due to the testing of samples in the omicron period characterized by the prevalence of a substantially mutated S RBD domain.

Our proof of principle study was limited in that the analytical sensitivities of the disease-specific commercial ELISAs were poor and wastewater-derive measurements semi-quantitative, at best. Additional work is needed to determine the best enrichment and detection methods that would reliably produce quantitative trends while mitigating potential matrix effects and preserving effector functions (e.g., epitope binding). To that end, we tried standard Ig enrichment methods employing protein A/G, L, as well as bulk protein concentration methods such as PEG and acetone but did not observe significant enhancement of anti-S IgA or IgG signals (data not shown). Moreover, we attempted to ascertain anti-N protein IgA but very low and non-titratable signal was observed with the commercial kit used, which suggested a non-specific signal component (data not shown).

### Applications to Public Health and epidemiology

Using the example of SARS-CoV-2 Spike protein, studies have consistently demonstrated that mucosal (saliva) antibody titres to this antigen that increase within the first 2 weeks following infection consist of a long-lived IgG (3+ months), and relatively short-lived (<2 months) IgA and IgM titres (25,62). In addition to acting as proxy markers of infection or vaccination, specific antibodies can be COPs. High serum IgA (non-polymeric) is associated with severe COVID-19 (63,64). However, the effector functions of IgA sub-classes vary between body compartments; for example between mucosal SIgA and serum IgA (60).

A key takeaway from our results is that the major proportion of wastewater-derived human antibodies, in particular IgA isotypes, are mucosal in origin. This is critical knowledge in our interpretation of the signal and what it might mean regarding the course of infection, and protection from disease in a population. Mucosal SARS-CoV-2 S IgA titres increase with SARS-CoV-2 infection (65) and negatively correlate with viral load/shedding in the nasal mucosa during the course of infection (66). Moreover, high levels of mucosal IgA, but not IgG are observed in re-infected vs. naïve cohorts, and this protects against breakthrough infections in vaccinated individuals (67).

How might this knowledge be applied and interpreted in the wastewater context? Increasing wastewater-based SARS-CoV-2 RNA signal should forecast increasing S protein SIgA within 2 weeks and the magnitude and latency of that response should be indicative of the degree of community immunity and viral transmissibility. A blunted and/or delayed response relative to S protein IgG might be indicative of poor immunity and increased transmission risk (due to reduced Ig secretion in the nasopharynx). Current intramuscular COVID-19 mRNA vaccines do not reliably induce a mucosal IgA response although there appears to be a prime-boost effect of vaccination subsequent to infection, and increased mucosal anti-S IgG is a more reliable indicator of vaccination (68). In the wastewater context, an anti-S IgG wave in the absence of accompanying anti-S IgA could be a reliable population-level proxy of intramuscular vaccine uptake. Applied at neighbourhood-scale, this information could be used to better target vaccine pop-up clinics and similar interventions. Wastewater-based anti-S IgA trends have the potential to serve as measures of efficacy for any future vaccines that are designed to induce mucosal immunity. A longitudinal survey of wastewater fractions probing for SARS-CoV-2 RNA as well as S protein IgA and IgG is needed to derive potential immunological thresholds for protection from infection or consequential disease.

This study points to a functional (antigen-binding) repertoire of SIgA and IgG present in wastewater solids. While preserved binding function was observed for IAV and SARS-CoV-2 antigens, it is highly likely that the repertoire recognizes a broader range of epitopes. Future studies may elaborate on these, identifying novel epitopes or correlates of human pathology. It is expected that further characterization of the wastewater-derived antibody repertoire might also spur discovery in aspects of non-human Igs.

Notwithstanding the need to establish ultrasensitive methods and sufficient baseline readings to effectively interpret wastewater-derived antibody signals, four major epidemiological and public health applications can be envisioned: 1) tracking the history of exposure to vaccine and non-vaccine antigen(s) in a population, 2) inferring population-level protection from infection or consequential disease, 3) monitoring the real-world efficacy of new vaccine formulations and/or campaigns, and 4) agile targeting of public health interventions (e.g., to a specific neighbourhood, building).

## MATERIALS AND METHODS

### Sample collection

Primary sludge and post-grit influent samples were collected from the single WRRF serving Ottawa (ROPEC; serving a population of ∼1M) and kept at 4°C for <72 hours before being processed fresh or archived/stored at -20°C. Additional influent samples were collected from buildings at the University of Ottawa campus, and residential neighbourhoods in the Ottawa region. Influent samples from emergency housing facilities in the downtown area of Toronto were collected twice a week and immediately shipped to Ottawa at 4°C where they were processed fresh or archived/stored at -20°C. Stool samples were collected from hospital in-patients admitted for COVID-19 and plasma was obtained from healthy volunteers. No identifying information was recorded for wastewater, stool, or plasma samples. The University of Ottawa research ethics board deemed that no ethics review was required for this study because no personal information on residents where wastewater was sampled was collected. The resulting concentrated liquid fractions and solid pellets were then used immediately or stored at -20 °C until further use. Stool samples (50-200 mg) were homogenized in 1X PBS containing protease inhibitors and 0.5% Triton X-100 then centrifuged for 10 minutes at 15000 x g to clarify the sample. Plasma was obtained by centrifuging heparin-containing blood samples for 10 minutes at 750 x g.

### Primary sludge processing

Solids from primary sludge (approximately 250 mg) were collected by centrifugation (15000 x *g*, 4 °C, 10 min), whilethe supernatant was retained as the liquid fraction and concentrated (∼30x) using filters with a molecular weight cut-off of 10 kDa (Amicon, UFC9010). A liquid fraction of the sludge was harvested by diluting 5 mL of primary sludge 1:10 with 1X PBS, pH 7.4, briefly mixing to homogenize, clarifying by centrifugation (3000 x *g*, 4°C, 10 min), filtering (0.45 µm), and then concentrating the supernatant (∼ 50x) using filters with a molecular weight cut-off of 10 kDa (Amicon, UFC9010).

### Influent processing

Solids from post-grit influent and influent samples (influent solids; approximately 100-300 mg) were collected by centrifugation (3000 x *g*, 4°C, 10 min). The supernatant was retained and concentrated (∼50x) using filters with a molecular weight cut-off of 10 kDa (Amicon, UFC9010). The pellet was resuspended in residual liquid and transferred to a new tube and then pelleted again by centrifugation (15000 x *g*, 4°C, 10 min). The small amount of supernatant was then removed by aspiration resulting in the final pellet.

### Protein extraction from solid material

Proteins were extracted by bead beating the wastewater solids obtained above in homogenization buffer (1x PBS, 0.5% Triton X-100, 1 mM PMSF, 1x Protease Inhibitor cocktail; MilliporeSigma 4693116001) for 3-4 cycles of 30 seconds at 4°C, 5.0 m/s using an OMNI Bead Ruptor 24. The resulting homogenate was separated by centrifugation (15000 x *g*, 4 °C, 10 min) and the supernatant was clarified by an additional centrifugation step (15000 x *g*, 4 °C, 5 minutes). After collection of the clarified supernatant (**Fig 1A**; sludge or influent solids), samples were analyzed immediately or stored at -20 °C until later use. All liquid fractions from either sludge or influent did not require homogenization and were assayed as raw concentrates (**Fig 1A**; sludge or influent liquid concentrates).

### Total Protein quantification

The concentration of total protein in each processed sample was quantified using the DC Protein Assay Kit II (Bio-Rad) with a standard curve (R^2^ > 0.99) ranging from 100 – 0.2 mg/mL of Bovine Serum Albumin (MilliporeSigma). Samples were assayed in technical triplicate and the resulting absorbance values at 750 nm were measured using the BioTek Synergy HTX Multimode Reader (Agilent). Readings were corrected by subtraction of the blank and converted to concentrations by extrapolating from the standard curve.

### ELISA

The amounts of immunoglobulins were determined using commercially available ELISA kits: Human IgG (Novus, NBP2-60474), Human IgA (Novus, NBP2-60509), Human IgM (Novus, NBP2-60477), Human IgA (Abbexa, abx151955), Human Secretory IgA (Abbexa, abx253169), Human Anti-Influenza virus A IgA (Abcam, ab108743), Human Anti-Influenza virus A IgG (Abcam, ab108745), Anti-Respiratory Syncytial Virus (RSV) IgA Human (Abcam, ab108764), Anti-Respiratory Syncytial Virus (RSV) IgG Human ELISA Kit (Abcam, ab108765), SARS-CoV-2 Spike Protein Serological IgG ELISA Kit (Cell Signaling, #20154C), and SARS-CoV-2 Spike Protein Serological IgA ELISA Kit (Cell Signaling, #58873C). Samples were tested with dilutions ranging from 1-125x, and with analyte volumes of 50-100 µL depending on the requirements of each specific assay kit. All samples were assayed in duplicate or triplicate and the resulting absorbance values at 450 nm were measured using the BioTek Synergy HTX Multimode Reader (Agilent). Readings were corrected by subtraction of the buffer blank, converted to concentrations by interpolating from the standard curve, and then scaled by the dilution factor.

### Stain-free SDS-PAGE

Polyacrylamide gels were made according to the manufacturer’s instructions using the TGX Stain-Free FastCast Acrylamide Kit, 10% (Bio-Rad) with TEMED (Bio-Rad) and ammonium persulfate (Bio-Rad). Prior to loading the gel, 4X Laemmli Sample Buffer containing 10% 2-Mercaptoethanol (Bio-Rad) was added to each sample to a final concentration of 1X. Samples were thoroughly mixed, heated at 95 °C for 5 min, and then clarified by centrifugation (15000 x *g*, 25°C, 3 min). Samples were loaded (20 µL) and run in Tris-glycine buffer (25 mM Tris-base, 192 mM glycine, 0.1% SDS) at 200 V for 45-90 minutes depending on the desired separation of the protein ladder (GeneDireX, PM008-0500). After electrophoresis, the gel was imaged (Bio-Rad, ChemiDoc Touch) with UV activation (45 seconds) to visualize total protein.

### Western Blot

Following SDS-PAGE, proteins were transferred to a nitrocellulose membrane (Bio-Rad) using a Tetra wet blotting Module (Bio-Rad) with Tris-glycine buffer (25 mM Tris-base, 192 mM glycine, 20% methanol) at 100 V for 90 minutes at 4°C. The membrane was then rinsed with distilled water and stained with Ponceau S solution (0.5% + 1% acetic acid) to visualize transfer efficiency of total protein. Excess stain was washed away with distilled water before the membrane was blocked with skim milk buffer (4% in 1X TBS + 0.1% Tween 20) for 30 minutes at 25°C. Monoclonal antibodies targeting Human IgA (Invitrogen, MA5-32575), IgG (Invitrogen, MA5-42729), polyclonal IgM (Invitrogen, PA5-86030), and J chain (MA1-80527) were diluted 1:1000 in the same blocking buffer. The initial blocking buffer was removed and replaced with solutions containing diluted antibodies. Membranes were incubated with primary antibodies overnight at 4 °C with gentle shaking. Primary antibody solutions were removed the next day, and the blot was washed in TBS-T (1X TBS + 0.1% Tween 20) for 5 minutes at 25°C with gentle shaking. This wash step was repeated 4 more times for a total of 5 washes, while replacing the wash buffer each time. The final wash buffer was removed, and a secondary HRP-linked antibody targeting rabbit IgG (Cell Signaling, 7074S) or mouse IgG (Cell Signaling, 7076S) diluted 1:3000 in blocking buffer was incubated with the blot for 30 minutes at 25°C with gentle shaking. The secondary antibody solution was then removed, and the wash steps were repeated as described above. The washed blot was then incubated with Clarity Western ECL Substrate (Bio-Rad, 1705060). Excess ECL substrate was removed, and blots were imaged using the Bio-Rad ChemiDoc Touch.

### Mass spectrometry (MS)

The same procedure was followed as outlined in the previous method sections, but with one exception: Solids were extracted using a buffer (1X PBS, 1 mM PMSF, 1X Protease Inhibitor cocktail) containing 0.5% of the MS-compatible detergent digitonin in place of Triton X-100. A liquid fraction of the sludge containing no detergent was harvested by diluting 5 mL of primary sludge 1:10 with 1X PBS, pH 7.4, briefly mixing to homogenize, clarifying by centrifugation (3000 x *g*, 4°C, 10 min), filtering (0.45 µm), and then concentrating the supernatant (∼ 50x) using filters with a molecular weight cut-off of 10 kDa (Amicon, UFC9010). Following SDS-PAGE, bands were excised from the gel and stored in water until being submitted for analysis. Sample work-up, processing, and analysis were completed by the uOttawa Proteomics Resource Center as a service. Intensity values for detected peptides were log-normalized to a relative scale and visualized as a heat map. Mapped protein IDs along with peptide numbers and intensity data are presented in Data_S4 (csv file).

## Supporting information

Data S4_mapped peptides

## ACKNOWLEDGEMENTS

We are thankful for the sample collection and shipping support provided by Nora Dannah, Karl Cruz, Alexandra Johnston, Claire Gibbs, and Wyatt Weatherson. Mass spectrometry services were provided by Dr. Zhibin Ning of the Proteomics Resource Centre at uOttawa.

