## Supplementary material for "Urban wastewater contains a functional human antibody repertoire of mucosal origin": Fig S1

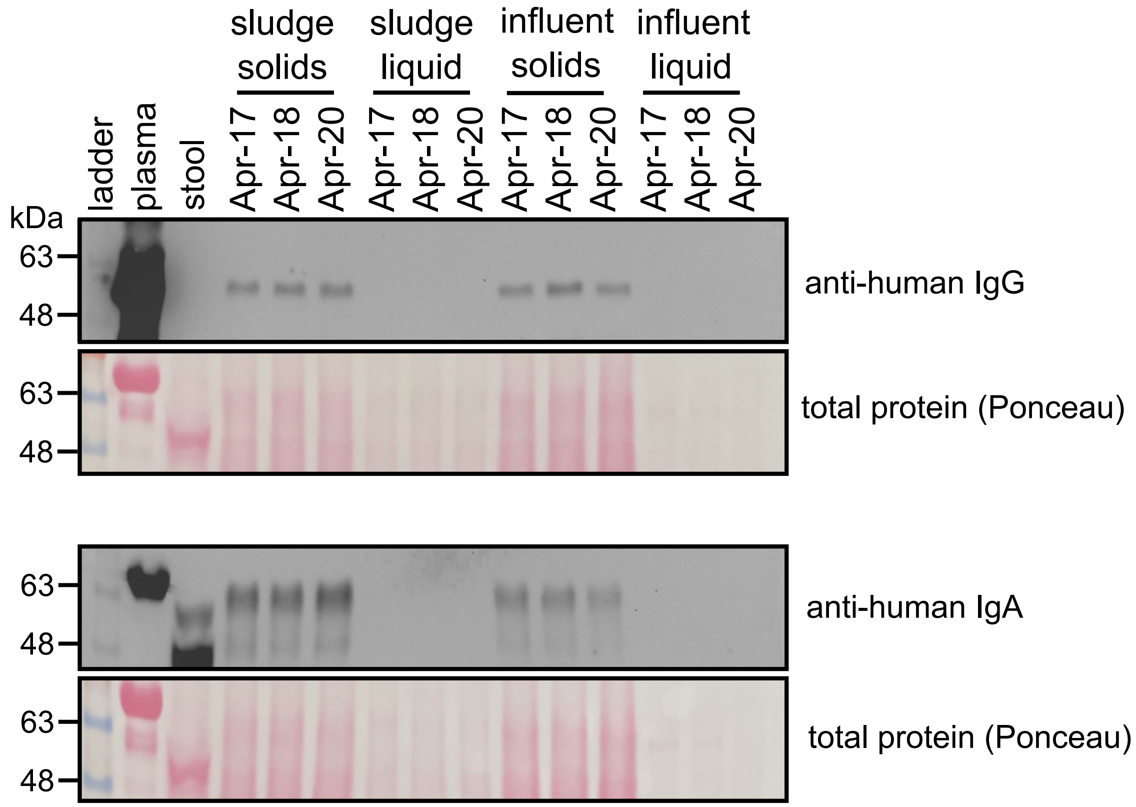


**Figure S1. Concentrated liquid fractions have a low soluble protein content.** Proteins from solids or concentrated liquid fractions processed from samples collected on three different dates were resolved by SDS-PAGE and transferred to a nitrocellulose membrane. Blots were probed with anti-human IgG (top) or IgA (bottom) and accompanying ponceau-stained membranes showing total protein loaded. Plasma and stool samples serve as specificity controls.
