## Supplementary material for "Urban wastewater contains a functional human antibody repertoire of mucosal origin": Fig S2

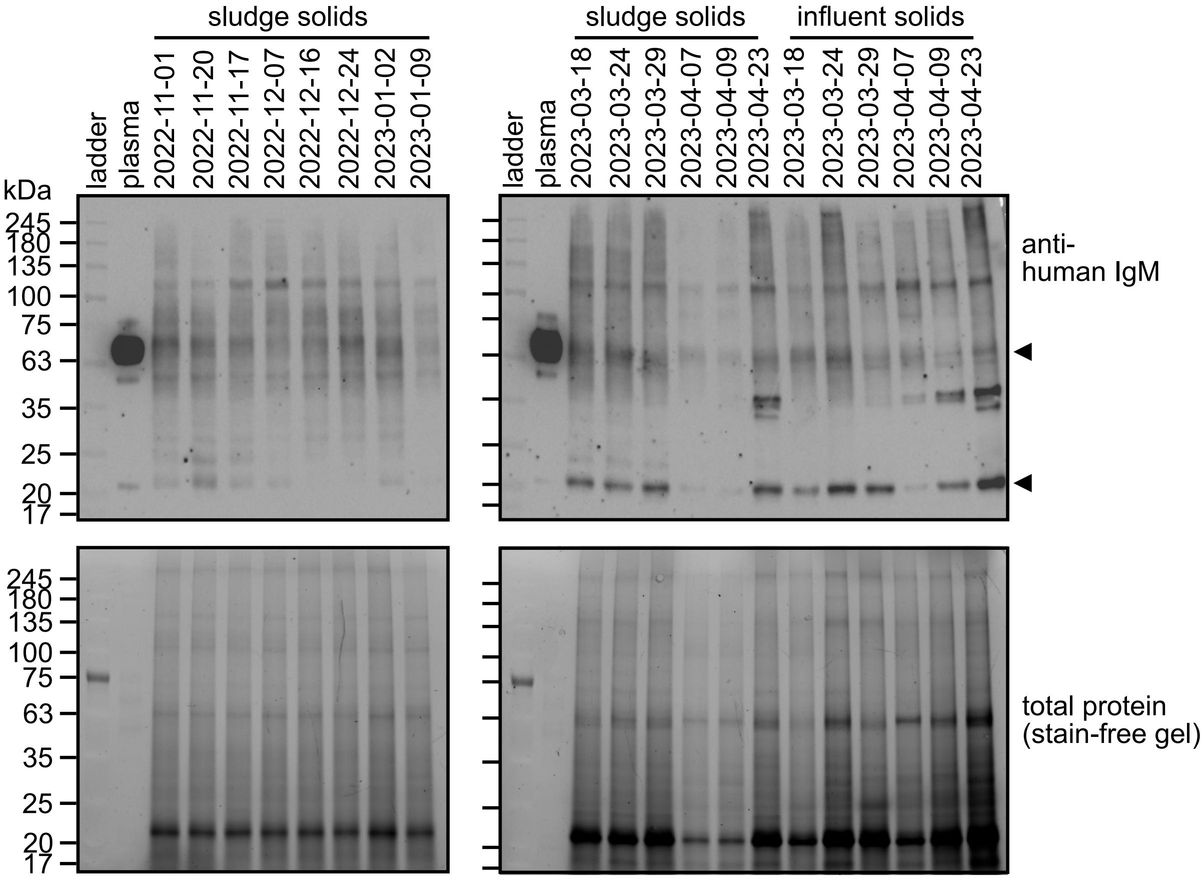


**Figure S2. Presence of IgM chains in wastewater solids.** Proteins from solids or concentrated liquid fractions processed from samples collected on different dates were resolved by SDS-PAGE and transferred to a membrane. Blots were probed with a polyclonal human antibody (top) and the accompanying stain-free gels (bottom) show total protein loaded. Plasma sample serves as a specificity controls. The top arrow indicates the the IgM heavy chains, with class-agnostic light chains indicated by a lower arrow, likely as a result of the polyclonal nature of the antibody.
