## Supplementary material for "Urban wastewater contains a functional human antibody repertoire of mucosal origin": Fig S3

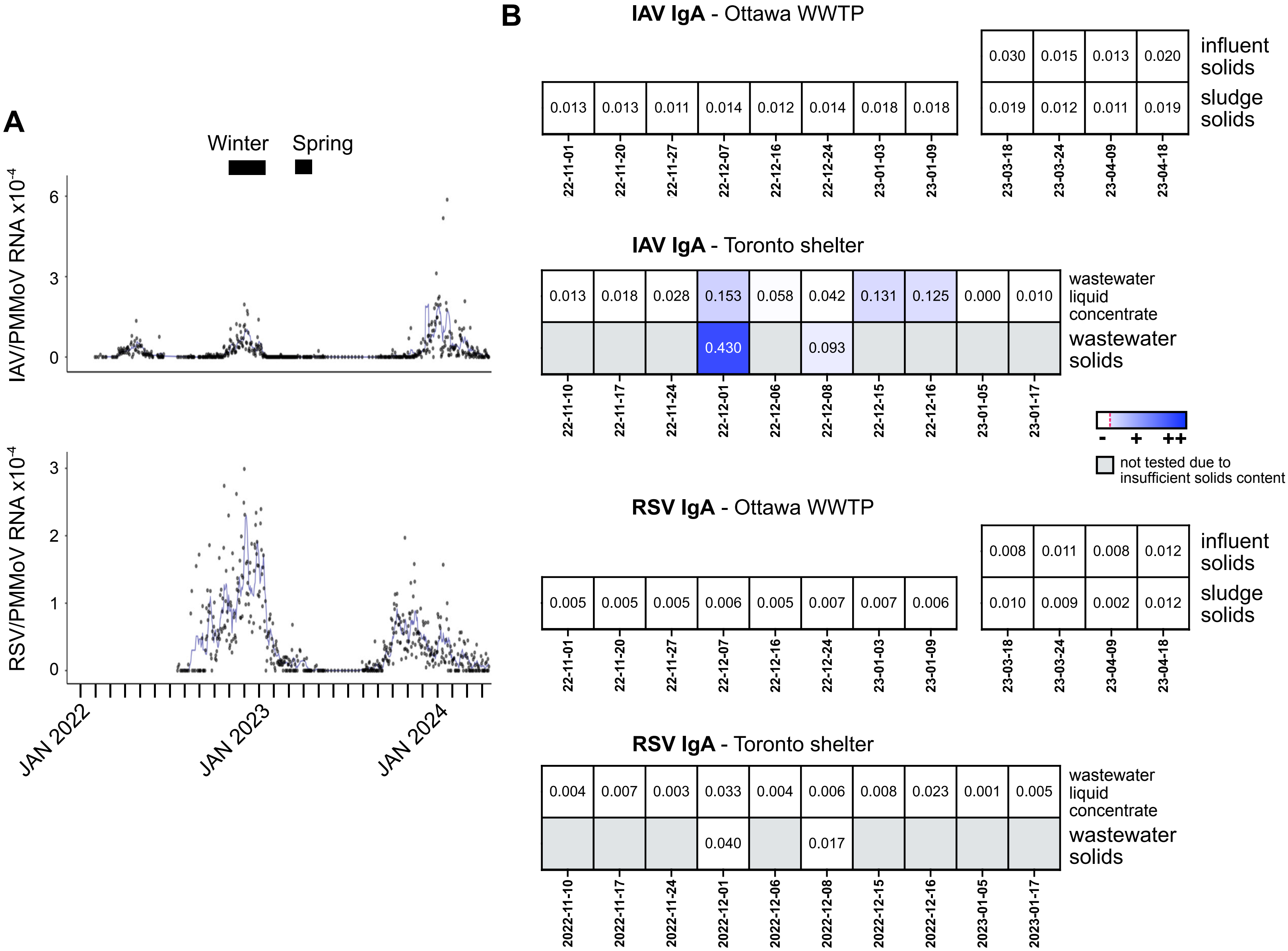


**Figure S3. Survey of anti-IAV and RSV IgA in WWTP and buidling-level wastewaters.** (**A**) Scatter plot of wastewater-based IAV and RSV RNA signals at Ottawa WWTP from January 2022 until April 2024. Data is normalized to PMMoV and polyline of 7-day moving average is plotted (blue). Ig analysis was perfomed on archived samples from Winter 2022 (high IAV/RSV incidence) and Spring 2023 (low incidence) periods collected from Ottawa WWTP and a Toronto shelter. (**B**) Direct ELISA detecting either anti-IAV or anti-RSV IgA in samples described in A. Values represent the average of technical duplicates and are corrected against a buffer blank.
